# Three Dimensions of Association Link Migraine Symptoms and Functional Connectivity

**DOI:** 10.1101/2021.03.31.437905

**Authors:** Samuel R. Krimmel, Danielle D. DeSouza, Michael L. Keaser, Bharati M. Sanjanwala, Robert P. Cowan, Martin A. Lindquist, Jennifer Haythornthwaite, David A. Seminowicz

## Abstract

Migraine is a heterogeneous disorder with variable symptoms and responsiveness to therapy. Due to previous analytic shortcomings, variance in migraine symptoms has been weakly and inconsistently related to brain function. Taking advantage of neural network organization measured through resting-state functional connectivity (RSFC) and advanced statistical analysis, sophisticated symptom-brain mapping can now be performed. In the current analysis we used data from two sites (n=102 and 41), and performed Canonical Correlation Analysis (CCA), relating RSFC with a broad range of migraine symptoms ranging from headache characteristics to sleep abnormalities. This identified three dimensions of covariance between symptoms and RSFC. Importantly, none of these dimensions bore any relationship with subject motion. The first dimension was related to headache intensity, headache frequency, pain catastrophizing, affect, sleep disturbances, and somatic abnormalities, and was associated with frontoparietal and dorsal attention network connectivity, both of which are major cognitive networks. Additionally, RSFC scores from this dimension – both the baseline value and the change from baseline to post-intervention – were associated with clinical responsiveness to mind-body therapy. The second dimension was related to an inverse association between pain and anxiety, and to default mode network connectivity. The final dimension was related to pain catastrophizing, and salience, sensorimotor and default mode network connectivity. These unique symptom/brain-mappings over three dimensions provide novel network targets to modify specific ensembles of symptoms. In addition to performing CCA, we evaluated the current clustering of migraine patients into episodic and chronic subtypes, and found no evidence to support this clustering. However, when using RSFC scores from the three significant dimensions, we identified a novel clustering of migraine patients into four biotypes with unique functional connectivity patterns. These findings provide new insight into individual variability in migraine, and could serve as the foundation for novel therapies that take advantage of migraine heterogeneity.

## Introduction

Migraine is a common, poorly managed, and disabling disorder, affecting over 10% of the global population (Leonardi & Raggi, 2013; Lipton et al., 2001, 2007; Rasmussen, 1995). Even with improved treatments (Edvinsson et al., 2018; Goadsby et al., 2017; Seminowicz et al., 2020; Yuan et al., 2019), it is clear that continued research is warranted. While its diagnosis is based on migraine attacks, migraine is a much more complex disorder, often featuring sleep, affective, cognitive, and general health abnormalities (Breslau et al., 1994, 2000; Karthik et al., 2012; Lin et al., 2016). Importantly, the presentation of these symptoms is heterogeneous across subjects, which is an important consideration for migraine research and treatment. Improved understanding of this variability would allow for precision medicine, and would also help to stratify migraine patients into possible subcategories.

The brain can be functionally segregated into structured patterns of covariance, called networks (Yeo et al., 2011), and these brain networks have been linked to migraine symptoms (Schwedt et al., 2015). A common method for identifying brain networks is resting state functional connectivity (RSFC), a measure of functional coupling between regions of the brain (McIntosh, 2000), typically estimated through correlating timeseries from functional units in functional magnetic resonance imaging (fMRI) acquired during rest (i.e., in the absence of a task or stimulus). These studies typically associate RSFC with individual migraine symptoms, but do not take into account that symptoms are interdependent, as are estimates of RSFC.

Canonical Correlation Analysis (CCA) (Hotelling, Harold, 1935) is a multivariate statistical approach that linearly transforms variables from two sets of data to maximize the correlation between them. CCA has become increasing popular in neuroimaging (H.-T. Wang et al., 2020) and has identified dimensions of association between brain connectivity and demographic information (Smith et al., 2015), psychopathology (Drysdale et al., 2017; Mihalik et al., 2019; Xia et al., 2018), and mind-wandering (H.-T. Wang et al., 2018). In the current study we sought to identify dimensions of covariance between a diverse range of migraine symptoms and RSFC. Using a large number of subjects from two research centers and CCA, we revealed three modes of association between migraine symptoms and RSFC. Unlike standard mass univariate analyses, these 3 dimensions were each informed by many clinical and RSFC features simultaneously. These dimensions related to symptoms globally and also to more specific combinations of symptoms, and reveal potential targets for brain stimulation or other interventions. Additionally, we found that one of these dimensions was associated with improvements from enhanced mindfulness-based stress reduction (MBSR+), an emerging non-pharmacologic therapy for chronic pain. Finally, we were able to show that the current clustering of migraine into episodic or chronic subtypes based on headache frequency alone is not supported, and propose a new data driven biotyping of migraine patients into 4 clusters that is based on RSFC and symptoms together.

## Methods

### Overview

We took clinical measures and RSFC from two study centers. 143 subjects had complete RSFC and clinical data, and were used as input data for the CCA. 166 subjects had complete clinical data and were used to assess the empirical support of episodic versus chronic subtypes of migraine. Multi-site harmonization was used to correct for effects of site For RSFC, while still maintaining clinical information. We first applied PCA to both clinical and RSFC data to address multicollinearity while reducing dimensionality.

Regularized CCA was then performed on components explaining 80% of variance and permutation tests were used to assess statistical significance. Canonical correlation is simply a correlation between canonical variates, where canonical variates are created from two domains of data (clinical and RSFC) so that the canonical correlation is maximized. Stated differently, CCA allows for dimensions of association to be found between clinical and RSFC, where these associations are formed through unique combinations of clinical and RSFC variables. Each canonical correlation consists of a clinical and RSFC canonical variate that are created to be associated with one another. For significant canonical correlations, RSFC canonical variates were then tested for association with motion (a major confound for RSFC studies) and clinical improvement from mind-body therapy. Finally, these scores underwent k-means clustering to identify biotypes in our sample (see Fig 1 for study overview).

**Fig 1.**
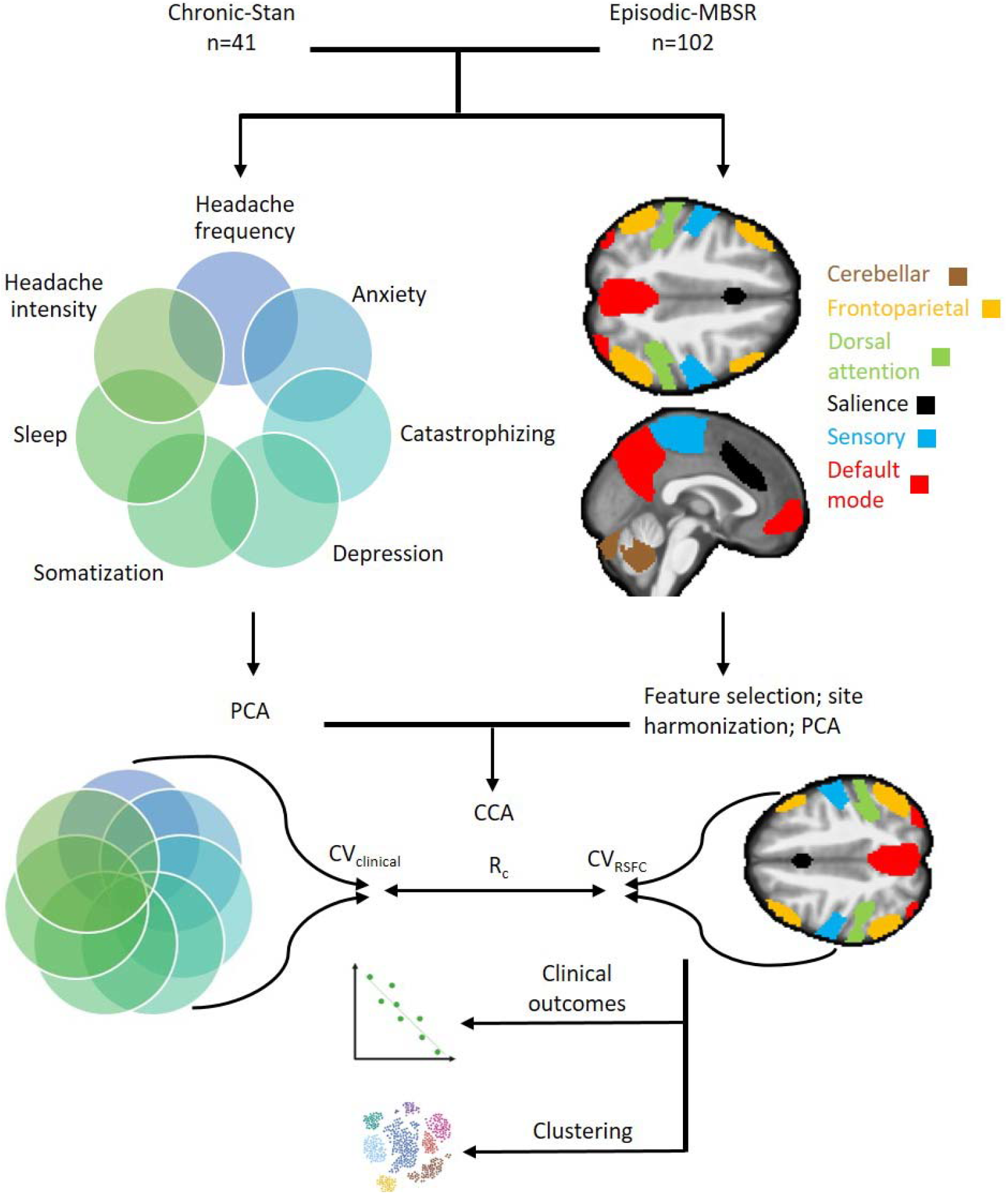
Study overview. For Canonical Correlation Analysis (CCA), 143 migraine subjects were collected from two sites. Seven clinical measures were acquired along with resting-state functional connectivity (RSFC) estimates from 24 regions spanning six functional networks. After site harmonization of RSFC and principal components analysis (PCA), CCA was used to associate clinical and biological data. CCA is used to study the relationship between variables from two domains of interest measured in one sample. In CCA, linear combinations of the input variables from both domains are created (called canonical variates) so that the correlation between the combined variables (canonical correlation, R_c_) is maximized. RSFC canonical variates were associated with clinical outcomes to mind-body therapy and underwent clustering analysis to reveal biotypes of migraine. PCA=principal components analysis; R_c_ =canonical correlation; CV_RSFC_ =canonical variates for RSFC; CV_clinical_ =canonical variates for clinical data.

### Participants

Data from two migraine neuroimaging studies were analyzed, referred to here as the *chronic-Stanford* and *episodic-UMB* datasets. In the *chronic-Stanford* dataset, participants were recruited to participate in research investigating biomarkers of chronic daily headache using clinical, behavioral, and magnetic resonance imaging (MRI) data at Stanford University. All subjects were over 18 years of age and met the International Classification of Headache Disorders ICHD-3 diagnostic criteria for chronic migraine as determined by a physician for a minimum of three months (International Headache Society, 2018). Patients were excluded if they had any MRI contraindications or a history of severe neurologic/psychiatric disorders. Subjects provided written informed consent in accordance with Stanford University guidelines. MRI scans used in this analysis were a structural T1-weighted scan (repetition time [TR] 5900 ms, echo time [TE] minimum, flip angle 15°, voxel size 0.9 x 0.9 x 1 mm), and an eyes closed resting state functional scan (8 minutes, gradient echo spiral-pulse, TR 2000, TE 30 ms, slice thickness 4 mm, FOV 220 mm, flip angle 80°, voxel size 3.4 x 3.4 x 4.5 mm). For analyses of clinical data only, subjects were excluded due to missing data, leaving a total of 46 subjects for analysis (average age of 39, s.d. = 13, 38 female). For MRI analyses, subjects were excluded due to poor MRI quality, excessive motion during resting state (average framewise displacement greater than 0.4 mm), and missing clinical data, leaving a total of 41 subjects for analysis (average age of 38, s.d. = 13, 34 women).

The second dataset, *episodic-UMB*, consisted of data from a previously described clinical trial assessing enhanced mindfulness-based stress reduction (MBSR+) treatment of migraine (Seminowicz et al., 2020). Briefly, participants were between the ages of 18-65 and met the ICHD-3 criteria for episodic migraine with or without aura for more than a year (International Headache Society, 2018). Participants were excluded if they had a history of mindfulness training, reported severe or unstable psychiatric symptoms, and/or used opioids. All patients provided written informed consent in accordance with University of Maryland Baltimore standards. MRI scans used in this analysis were a structural T1-weighted scan (TR 2300ms, TE 2.98 ms, slice thickness 1 mm, FOV 256 mm, flip angle 9°, and voxel size 1 × 1 × 1 mm) and an eyes open resting state scan (10 minutes, echo planar imaging, TR 2000 ms, TE 28 ms, slice thickness 4 mm, FOV 220 mm, flip angle 77°, and voxel size 3.4 × 3.4 × 4 mm). For analyses of clinical data only, subjects were excluded due to missing data, leaving a total of 120 subjects for analysis (average age of 38, s.d. = 12, 106 female). For analyses of MRI data, subjects were excluded due to poor MRI quality, excessive motion during resting state (average framewise displacement greater than 0.4 mm), and missing clinical data, for a total of 102 subjects (average age of 38, s.d. = 12, 91 women). 83 of these subjects were also randomized to a clinical trial receiving either MBSR+ (n=42) or an active control condition (stress management for headache; n=41). These randomized participants received additional scans 10 and 20 weeks after the baseline scan (Seminowicz et al., 2020). Two MBSR+ subjects were excluded when analyzing changes in RSFC canonical variate scores leaving 40 subjects.

### Magnetic Resonance Imaging Preprocessing

Preprocessing was performed using FMRIPREP version stable (Esteban et al., 2020), a Nipype (Gorgolewski et al., 2017) based tool. Each T1-weighted (T1w) volume was corrected for INU (intensity non-uniformity) using N4BiasFieldCorrection v2.1.0 (Tustison et al., 2010) and skull-stripped using antsBrainExtraction.sh v2.1.0 (using the OASIS template). Spatial normalization to the ICBM 152 Nonlinear Asymmetrical template version 2009c (Fonov et al., 2009) was performed through nonlinear registration with the antsRegistration tool of ANTs v2.1.0 (Avants et al., 2008), using brain-extracted versions of both T1w volume and template. Brain tissue segmentation of cerebrospinal fluid (CSF), white-matter (WM) and gray-matter (GM) was performed on the brain-extracted T1w using fast (FSL v5.0.9 (Zhang et al., 2001)).

Functional data was slice time corrected using 3dTshift from AFNI v16.2.07 (Cox, 1996) and motion corrected using mcflirt (FSL v5.0.9 (Jenkinson et al., 2002)). This was followed by co-registration to the corresponding T1w using boundary-based registration (Greve & Fischl, 2009) with six degrees of freedom, using flirt (FSL). Motion correcting transformations, BOLD-to-T1w transformation and T1w-to-template (MNI) warp were concatenated and applied in a single step using antsApplyTransforms (ANTs v2.1.0) using Lanczos interpolation.

Frame-wise displacement (Power et al., 2014) was calculated for each functional run using the implementation of Nipype. ICA-based Automatic Removal Of Motion Artifacts (AROMA) was used to generate aggressive noise regressors as well as to create a variant of data that is non-aggressively denoised (Pruim et al., 2015). Many internal operations of FMRIPREP use Nilearn (Abraham et al., 2014), principally within the BOLD-processing workflow. For more details of the pipeline see https://fmriprep.readthedocs.io/en/stable/workflows.html.

### Resting State Functional Connectivity Processing

Resting-state denoising and seed-based analysis were conducted in the CONN functional connectivity toolbox version 19f (Whitfield-Gabrieli & Nieto-Castanon, 2012; http://www.nitrc.org/projects/conn). Given sustained controversy concerning global signal, we elected not to remove global signal in our analyses (Murphy & Fox, 2017). We performed aCompCor (Behzadi et al., 2007; Muschelli et al., 2014) to remove noise captured in white matter and CSF. We calculated the first five eigenvectors from four erosions of subject specific white matter masks and two erosions of subject specific CSF masks, as these levels of erosion no longer contain global signal (Power et al., 2017). To avoid noise being reintroduced while removing frequency specific noise, voxel-wise and regressor data were simultaneously band-pass filtered (0.008-0.15 Hz) and all data underwent linear detrending.

Following resting state functional denoising, timeseries were extracted from 32 ROIs based on an existing independent component analysis of the Human Connectome Project available from the CONN functional connectivity toolbox. These ROIs were organized into default mode, sensorimotor, visual, salience/cingulo-opercular, dorsal attention, frontoparietal, language and cerebellar networks. To estimate functional connectivity, the 32 ROI timeseries were correlated and the resulting Pearson correlation coefficient were converted to z scores. We excluded functional connections with visual and language network seeds, as we did not believe that these would be associated with the clinical symptoms in our analysis. Language network regions included parts of the posterior superior temporal gyrus and the inferior frontal gyrus (consistent with Wernicke’s and Broca’s area). Visual network regions included nodes in the occipital cortex. While the occipital cortex has been implicated in migraine, it is often associated with symptoms of aura (Hadjikhani et al., 2001), a symptom domain that was not measured in our two datasets. This left a total of 24 ROIs with 276 unique functional connections.

Study site has been shown to influence fMRI data and should be adjusted for when possible (Noble et al., 2017; Van Horn & Toga, 2009; Yu et al., 2018). To adjust for effects of site while controlling for clinical variables, age, and sex, we used Combining Batches of microarray data (ComBat) (Johnson et al., 2007). ComBat was developed to deal with batch effects in high dimensional data and uses an empirical Bayes method to remove batch effects. ComBat was applied following removal of irrelevant ROIs to create site corrected RSFC, which was used in all analyses except for when determining changes in canonical variate scores with therapy (see below section: *Canonical Variate Association with Clinical Outcomes of Treatment*).

### Clinical Features

Participants completed multiple self-report questionnaires to assess clinical characteristics. The two headache measures acquired were the frequency of headaches and the average pain associated with headaches. We also examined the exaggerated mental set applied to pain or anticipation of pain using the Pain Catastrophizing Scale (PCS; Sullivan et al., 1995). The Pittsburg Sleep Quality Index (PSQI) was used to measure sleep quality as well as sleep disturbances (Buysse et al., 1989). We used the Patient health questionnaire 15 (PHQ-15) to quantify somatization and somatic symptom severity (Kroenke et al., 2002). Finally, we examined affective health using the GAD-7 measure of anxiety (Spitzer et al., 2006) and the patient health questionnaire 9 (PHQ-9) measure of depression (Kroenke et al., 2001). The total score for each questionnaire (where applicable) was used for all analyses.

### Principal Components Analysis

We used Principal Components Analysis (PCA) to address multicollinearity and to reduce the dimensionality of our data. PCA creates a set of orthogonal components that can be linearly combined to explain all of the variance in the original dataset. We used singular value decomposition and selected the first 4 clinical components (Fig 2A), explaining more than 80 percent of the clinical variance, and the first 35 RSFC components, explaining more than 80 percent of the RSFC variance (Fig 2B). These components were used as input data for the canonical correlation analysis (see below section: *Regularized Canonical Correlation Analysis*).

**Fig 2.**
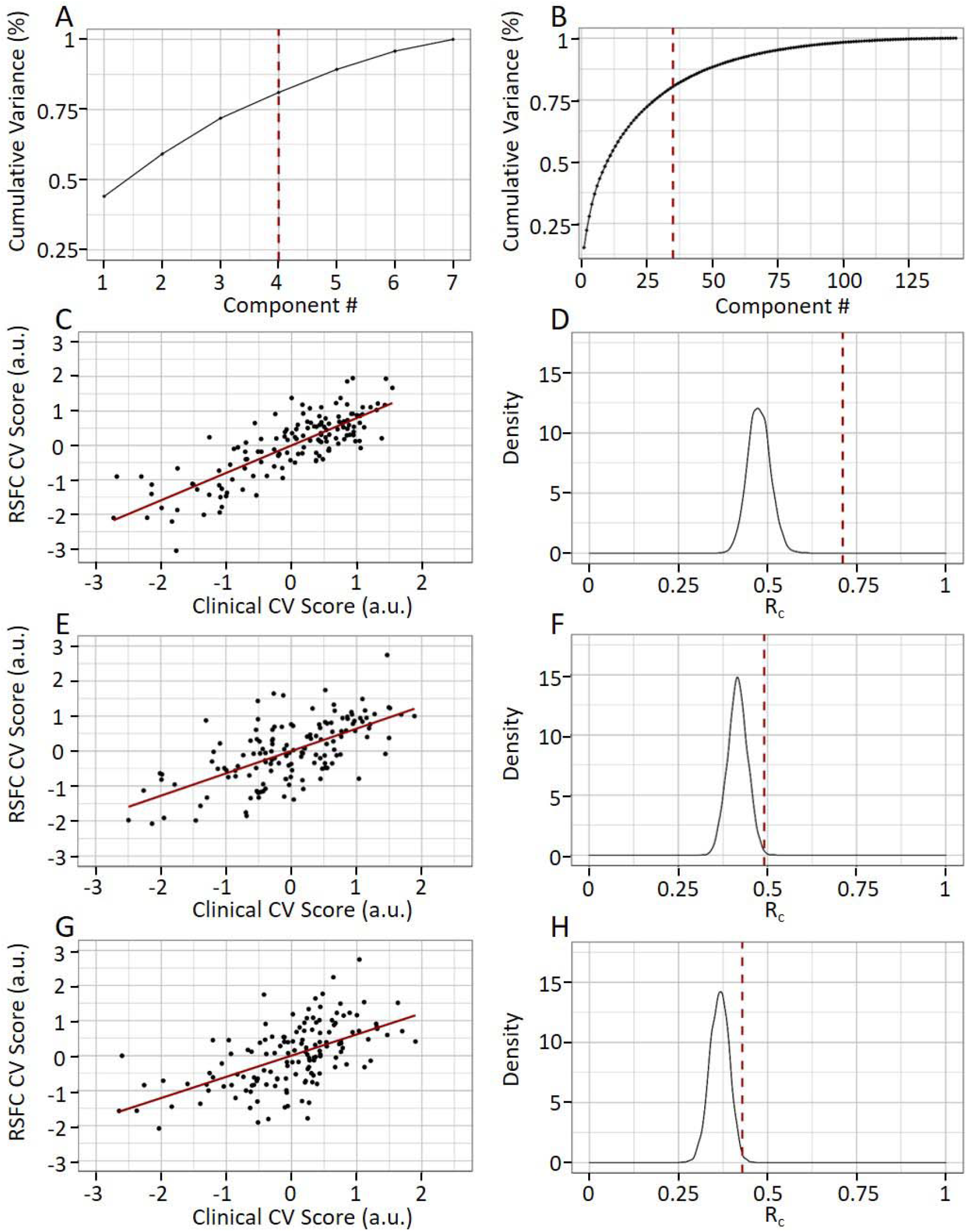
Using CCA to reveal three dimensions of association between clinical features and RSFC. Cumulative variance explained from clinical (A) and RSFC (B) principal components. Red lines indicate 80% variance explained. Canonical correlation (R_c_) between RSFC and clinical canonical variates for first (C), second (E), and third (G) canonical correlations. Permutation test for significance of R_c_ value (shown in red) relative to 5000 permutations for first (D), second (F), and third (H) canonical correlations. R_c_=canonical correlation; CV=canonical variate.

### Regularized Canonical Correlation Analysis

CCA is a method for identifying multivariate relationships between sets of variables (Hotelling, Harold, 1935). It is used to study the relationship between variables from two domains of interest (e.g., clinical symptoms and biological data) measured on the same set of individuals. In CCA, linear combinations of the input variables from both domains are created (called canonical variates) so that the correlation between the combined variables (canonical correlation, R_c_) is maximized. Similar to PCA it is possible to continue to seek additional pairs of canonical variates that are orthogonal to the preceding canonical variate(s). For each additional canonical variate pair the corresponding canonical correlation R_c_ decreases. The total number of possible pairs of canonical variates is given by the smaller of the number of variables in each domain, which in our analysis equaled four. CCA assumes minimal multicollinearity and is prone to overfitting, especially when the ratio of subjects to input variables is small. Previous attempts to associate clinical features with RSFC using CCA (Drysdale et al., 2017) may have been severely overfit (Dinga et al., 2019). In our analysis the starting subject to variable ratio was 0.5, which is likely to overfit, meaning that our results would be unlikely to generalize. Using PCA (described above), we reduced the subject to variable ratio to 3.4. While improved, this ratio was still worse than the generally recommended ratio of 10:1 (Pituch & Stevens, 2015; Tabachnick et al., 2007). To mitigate overfitting, we used an L2 norm regularized CCA (Gonzalez et al., 2008), and optimized regularization parameters (lambda) for clinical and RSFC inputs based on held out testing data from 10-fold cross-validation stratified for study site. The CCA model was built using 9 folds and the resultant canonical weights were used to create canonical variates in the held out fold. We repeated this process 10 times and for each repeat a new set of 10 folds was created, making in total 100 training sets with unique canonical weights and 100 corresponding testing sets to evaluate performance. For our final model, we used the regularization parameters that yielded the best average R_c_ value in held out testing data for the first canonical correlation.

### Reconstructing Model Weights

CCA was performed following PCA of clinical and RSFC data. However, the interpretation of the rotation matrix is challenging, and we therefore transformed the CCA weights back into their original space using the CCA model weights and the rotation matrix. Following this transformation, we were able to use the CCA model weights in the original clinical and RSFC spaces. Applying these weights to clinical and RSFC data yielded clinical and RSFC canonical variates respectively. We did not use these reconstructed weights directly for interpretation, and instead used canonical loadings for the purposes of interpretation (see below section: *Statistics* and figure 4). We did however use these weights to extract canonical variates over time.

### Canonical Variate Association with Clinical Outcomes of Treatment

We examined if the canonical variates obtained from CCA were associated with outcomes to MBSR+ in the *episodic-UMB* dataset. To do this, we examined subjects randomized to the enhanced mindfulness-based stress reduction condition and correlated baseline canonical variates for RSFC with the reduction in headache frequency after 20 weeks, the primary outcome of the trial (Seminowicz et al., 2020). We also sought to determine if changes in canonical variates were associated with reductions in headache frequency. To do this, we calculated the canonical variates using model weights from the CCA model described above, and calculated difference scores before and after therapy (20-week gap in between scans). In this analysis we did not use ComBat corrected RSFC data when determining changes in canonical variate scores, since the data were from a single site. The relationship between the change in canonical variate scores and reductions in headache frequency was assessed by means of correlation analysis over the corresponding time period.

### Determining Support for Episodic vs Chronic Migraine Clustering on Migraine Symptoms

Chronic versus episodic migraine can be thought of as a clustering solution to migraine and is defined based on a 15 headache day per month frequency cutoff. We sought to determine how well this clustering solution was supported by a more holistic catalogue of clinical data. We therefore determined the silhouette index for episodic versus chronic distinction using seven clinical variables in the combined sample (n=166). Should the episodic/chronic clustering have a silhouette index no better than chance (i.e., likely coming from a multivariate normal distribution), then is would argue against the current field norm of episodic and chronic migraine subtypes (see below section: *Statistics*).

### Data Driven Clustering of RSFC Canonical Variates

We sought to identify biotypes in migraine based on the significant RSFC canonical variates from CCA using k-means clustering. We used Euclidean distance as our distance metric for all clustering. We compared 26 clustering metrics using the NBclust package (Charrad et al., 2014) with cluster number (k) ranging from 2 to 10. For each value of k, a score for all 26 clustering metrics was computed, and cluster bperformance was ranked per metric (e.g., the value of k with the largest silhouette index received a rank of one). Per metric, a value of k ‘won’ if it received the first rank and therefore had the best performance for that specific clustering metric. We chose a final value for k using a majority rule, where the cluster number that had the most total victories (came in first for a clustering metric) was deemed the best overall. We paid special attention to the silhouette index, as this metric has been shown to accurately identify clusters of data (Petrovic, 2006).

### Statistics

To determine statistical significance of our CCA we performed a permutation test by constructing 5000 bootstraps with replacement (Efron, 2000) using the final model lambda parameters (see Results section: *Three Modes of Association Were Found Between RSFC and Migraine* Symptoms). For each bootstrap, we calculated R_c_ values for four canonical correlations and then compared this null distribution against true R_c_ values using a one-sided t-test for each of the four canonical correlations in a process previously described (Dinga et al., 2019). To interpret CCA results we calculated canonical loading by correlating canonical variates with their respective model inputs (e.g., RSFC values). We used a false discovery rate multiple comparisons correction on the resultant p values (Benjamini et al., 2001). Determining significance for clustering analysis can be challenging (Dinga et al., 2019; Liu et al., 2008). For our clustering, we tested the true silhouette index scores for the final clustering solution relative to 5000 multivariate Gaussian distributions based on the mean and covariance of the input data using a one sample t-test, in a process similar to ones previously described (Dinga et al., 2019; Liu et al., 2008). This approach allows us to test against the null model of a multivariate normal distribution which does not support clustering. A Spearman rank correlation was used for all correlation analysis unless stated otherwise. For all analyses, significance was determined using a 0.05 FDR-corrected threshold.

## Results

### Three Modes of Association Were Found Between RSFC and Migraine Symptoms

We used CCA to identify dimensions of covariance between a diverse array of clinical symptoms and RSFC. We optimized lambda values for regularization using performance in held out testing data to maximize canonical correlation (R_c_) for the first pair of canonical variates. The best performance in testing data for the first canonical correlation was an average R_c_ = 0.6 with lambda values of 0.3 for clinical data and 0.9 for RSFC, and we used these regularization parameters for our final model. In the final model, the first canonical correlation was R_c_ = 0.71, N = 143, p = 0.0002 (Fig 2C, D), the second was R_c_ = 0.49, N = 143, p = 0.0023 (Fig 1E, F), the third was R_c_ = 0.43, N = 143, p = 0.006 (Fig 1G, H), and the fourth was R_c_ = 0.35, N = 143, p = 0.098. The average R_c_ values for held-out testing data is as follows: first canonical correlation R_c_ = 0.6, s.d. = 0.19, second R_c_ = 0.14, s.d. = 0.25, third R_c_ 0.19, s.d. = 0.23, and fourth R_c_ = 0.03, s.d. = 0.23. We focused interpretation and further analysis on the first three significant canonical correlations. Note that each canonical correlation is between a pair of normally distributed clinical and RSFC canonical variates. Therefore, from the three significant canonical correlations, each subject has 3 RSFC and 3 clinical canonical variates.

### Association of RSFC Canonical Variates and Scanner Motion

Given that motion has profound effects on estimates of RSFC (Power et al., 2012), we ensured that the canonical variates for RSFC were unrelated to motion. We did not find a significant correlation between average framewise displacement during the resting state scan with the first r_s_ = 0.12, N = 143, p = 0.14 (Fig 3A), second r_s_ = 0.08, N = 143, p = 0.3 (Fig 3B), nor third r_s_ = −0.04, N = 143, p = 0.6 (Fig 3C) RSFC canonical variates. We therefore concluded that there was no evidence for a relationship between RSFC canonical variates and subject movement.

**Fig 3.**
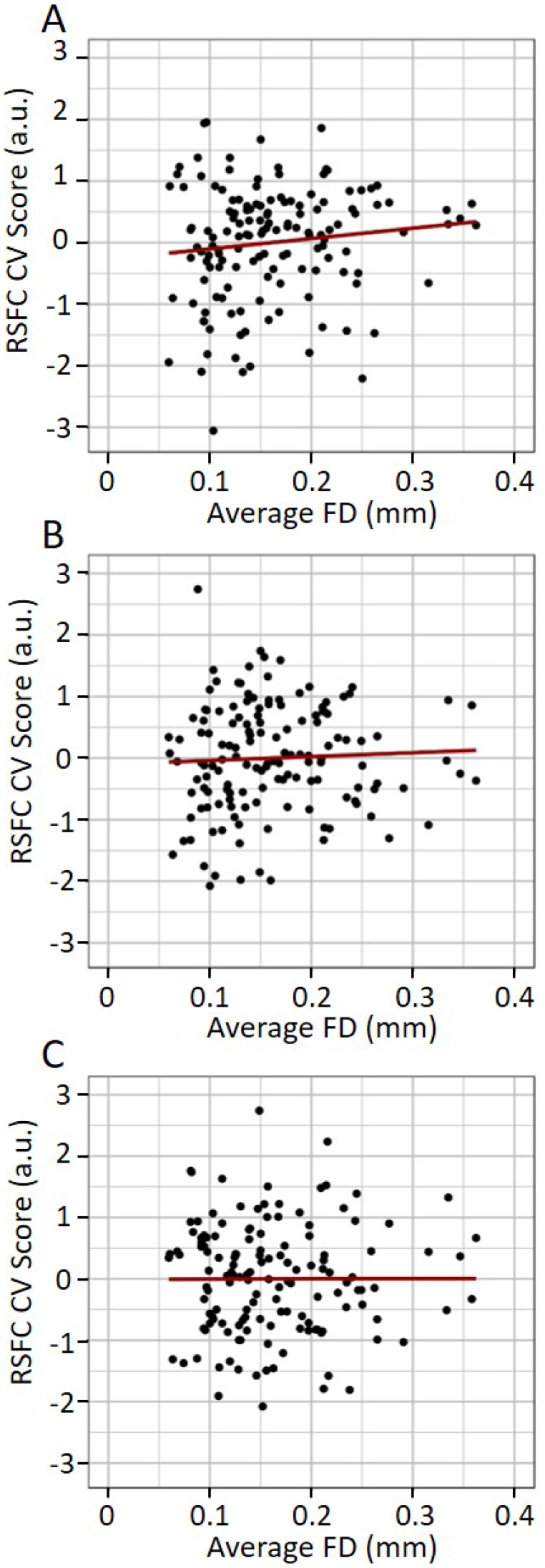
No association between canonical variates and subject motion. Scatter plot for first (A), second (B), and third (C) RSFC canonical variate relationship with average framewise displacement. CV=canonical variate, FD=framewise displacement.

### Interpretation of CCA Model

We correlated canonical variates with their respective RSFC/clinical symptoms (i.e., canonical loadings) to interpret the CCA model. While each of the three resultant dimensions consisted of clinical and RSFC canonical variates, we chose to name each dimension with a clinically derived name for ease of discussion. The first canonical variate for symptoms correlated in the same direction with all 7 clinical features (Fig 4A). Hence, we interpret the first clinical canonical correlation as the ‘global symptom dimension’. Patients with positive canonical variate values for the first canonical correlation had globally better symptoms (less severe symptoms overall). The associated first RSFC canonical variate primarily reflected frontoparietal network and dorsal attention network connectivity (Fig 4B).

**Fig 4.**
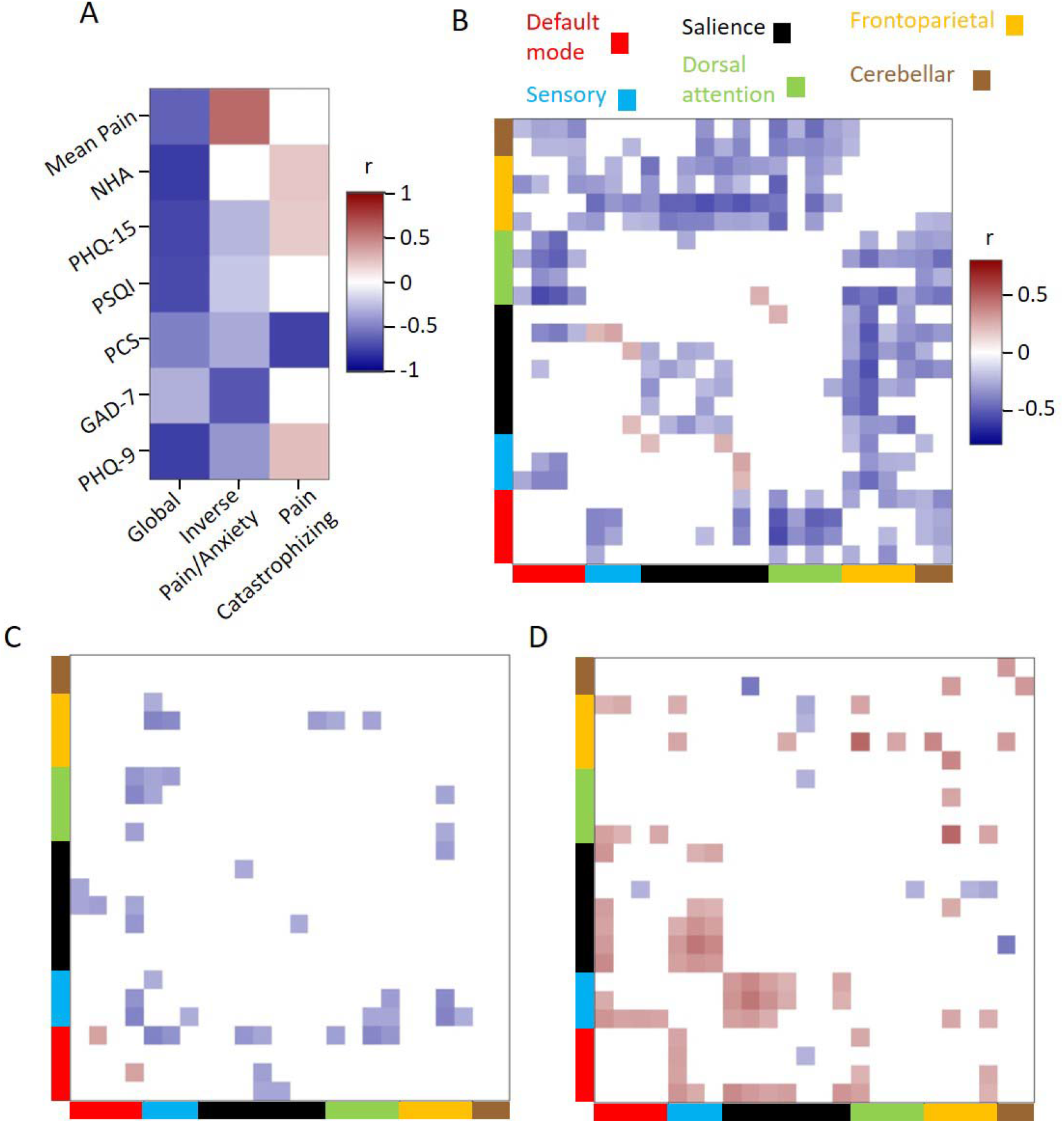
Interpretation of canonical variates. Canonical variates were correlated with input data to facilitate interpretation. A) For clinical data, correlations shown for the three identified dimensions. Correlation between RSFC and global dimension (B), inverse pain/anxiety dimension(C), and pain catastrophizing dimension (D) scores organized over 24 regions of interest in 6 networks. The global dimension was most associated with frontoparietal network and dorsal attention network connectivity with other networks. The inverse pain/anxiety dimension was most related to default mode network and sensorimotor connectivity. The pain catastrophizing dimension was most related to salience, sensorimotor and default mode network connectivity. The same color coding for networks and correlation color range is used in panels B, C, and D. All off-white correlation values are significant for all four panels. Mean Pain=average headache pain; NHA=headache frequency; PHQ-15=patient healthy questionnaire 15 item; PSQI=Pittsburgh sleep quality index; PCS=pain catastrophizing scale; GAD-7=generalized anxiety disorder; PHQ-9=patient health questionnaire 9 item; r=Pearson correlation coefficient.

The second clinical canonical variate reflected an inverse relationship between average headache pain and anxiety primarily, referred to here as the ‘inverse pain/anxiety dimension’ (Fig 4A). Patients with positive values had above average headache pain and below average anxiety. The pattern of functional connectivity was quite sparse for this but was primarily related to anti-correlation between the DMN and other networks (Fig 4C). The third symptom canonical variate primarily related to pain catastrophizing (Fig 4A) and related to salience, sensorimotor and default mode connectivity with one another (Fig 4 D). Patients with positive values had below average pain catastrophizing. We refer to this here as a ‘pain catastrophizing dimension’.

### RSFC Canonical Variates are Associated with Headache Reduction in MBSR+

We sought to determine if RSFC canonical variates were associated with improvements from mindfulness therapy for migraine. We correlated baseline canonical variates from functional connectivity with the reduction in frequency of headaches and found a significant association for the global dimension r_s_ = −0.44, N = 42, p = 0.0038 (Fig 5). RSFC canonical variates from neither the inverse anxiety/pain nor the pain catastrophizing dimensions were significantly associated with clinical improvement. Given that baseline global dimension RSFC canonical variate associated with clinical improvements, we next examined if changes in this dimension associated with headache frequency reduction following MBSR+ in the *episodic-UMB* cohort and found a significant association, r_s_= −0.40, N = 40, p = 0.014.

**Fig 5.**
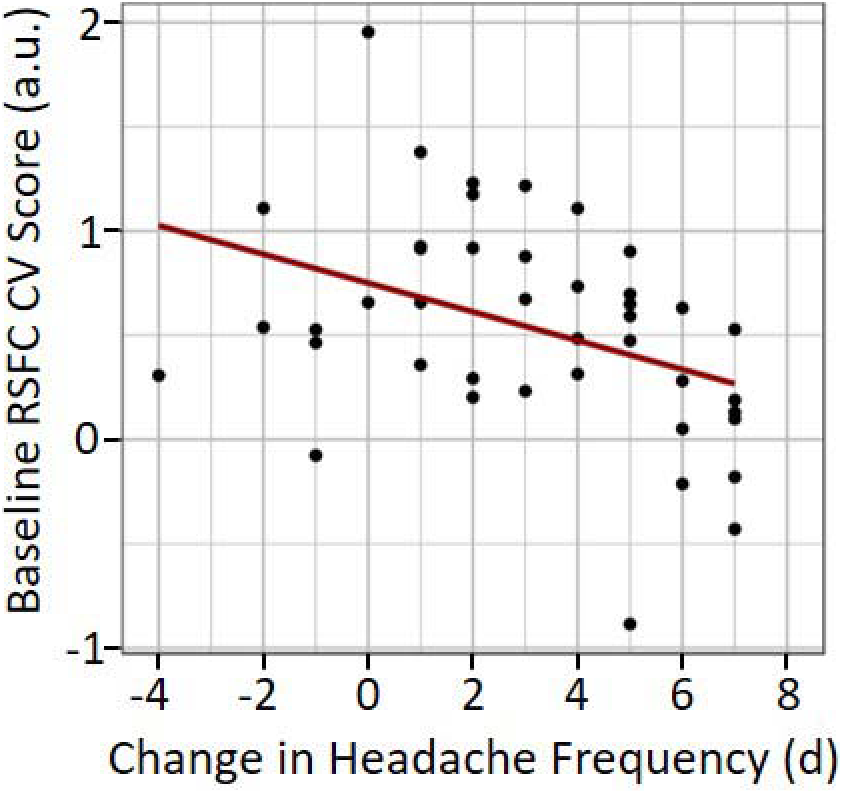
Association between baseline (pre-intervention) global symptom RSFC canonical variate and change in headache frequency after therapy (positive numbers indicate a reduction in headache days). Subjects with lower scores and more severe symptoms overall had better responses to MBSR+ than subjects were less severe symptoms. RSFC=resting-state functional connectivity; CV=canonical variate; d=days.

### Testing the Support for Current Migraine Clustering

We tested if the current clustering of migraine patients into episodic (<15 headaches/month) and chronic (≥15 headaches/month) subtypes differs from a multivariate Gaussian distribution. Chronic and episodic migraine patients were collected at unique sites, creating a potential confound (see discussion). Using the seven clinical features used for CCA, we determined the silhouette index in 166 patients using the chronic/episodic distinction, yielding a value of 0.23. We then again compared this to 5000 multivariate Gaussian distribution based on the mean and covariance of the 7 clinical measures. This analysis indicated that the chronic versus episodic clustering was not significantly different from a multivariate Gaussian distribution, N = 166, p = 0.66 (Fig 6A). Therefore, there is insufficient evidence to reject the null hypothesis that the episodic versus chronic clustering solution comes from a multivariate Gaussian distribution. This means that using a holistic catalogue of symptoms, we were unable to find support for subtyping migraine patients into episodic and chronic migraine categories.

**Fig 6.**
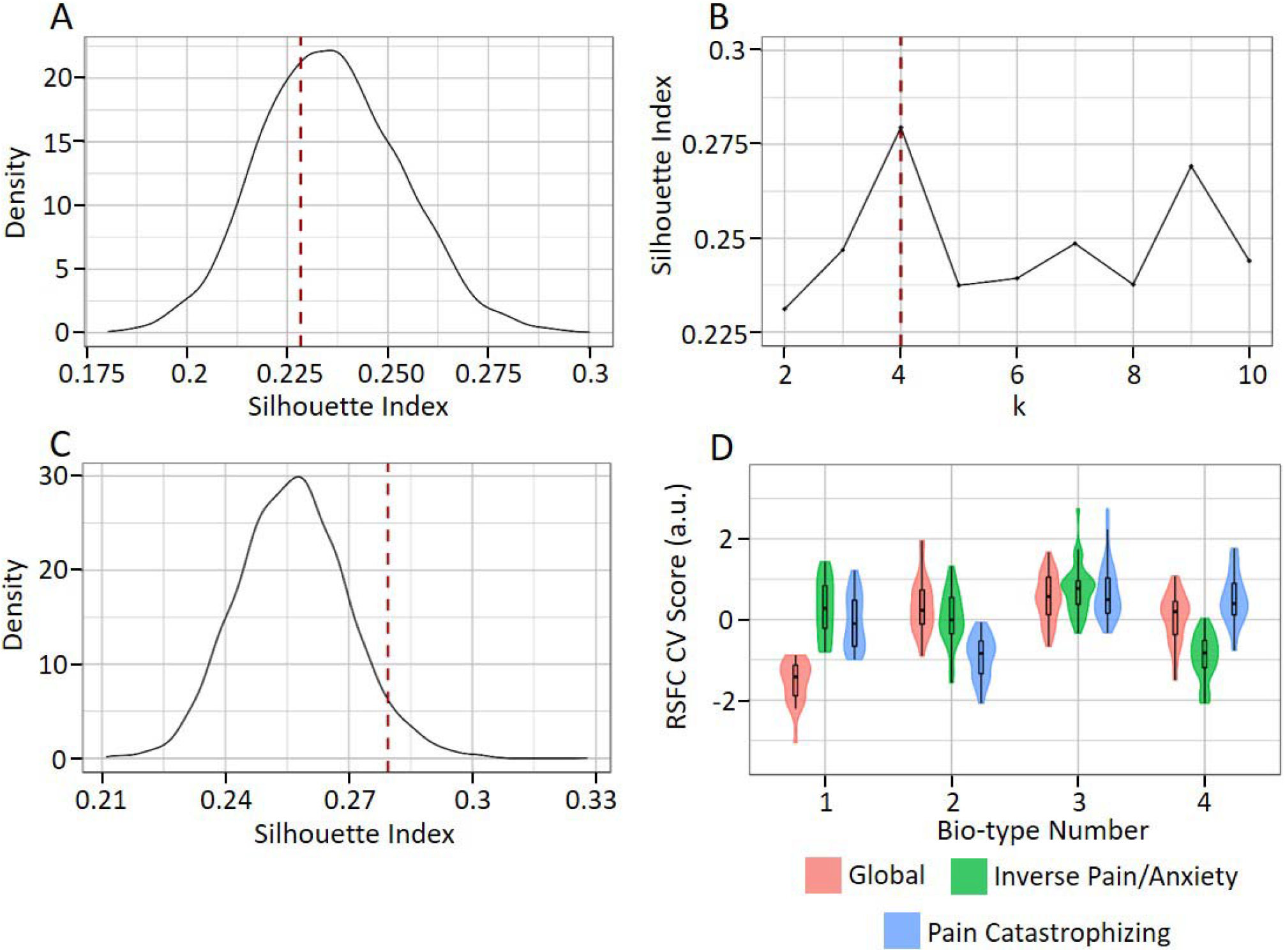
Lack of support for episodic/chronic migraine clustering and identification of a 4 biotype clustering of migraine. A) Lack of significance for clustering migraine into chronic and episodic subtypes based on silhouette index over 5000 multivariate Gaussian distributions of clinical data. B) Silhouette index for various cluster numbers (k) from three RSFC canonical variates indicating that k=4 had the best performance. C) Significance of clustering into four biotypes based on silhouette index over 5000 multivariate Gaussian distributions of RSFC canonical variate scores. D) Patterns of RSFC canonical variates for three dimensions over the four identified biotypes. Color coding indicates the dimension. Canonical variates are distributed with a mean of zero, therefore negative values indicate below average scores and positive values indicate above average scores. RSFC=resting-state functional connectivity; CV=canonical variate; k=number of clusters.

### Identifying Biotypes in Migraine

In order to identify subtypes of migraine based on biological data, called biotypes, we performed k-means clustering based on the canonical variates for RSFC for the three dimensions. The best performing solution was k=4, which had the best performance in 27% of metrics, though there were other solutions with similar performance (k=2 was best in 23%, no other solutions were consistently as strong; see Fig 6B for silhouette index scores over various values of k). To determine significance of a 4 cluster solution, we estimated 5000 multivariate Gaussian distributions based on the mean and covariance of the three RSFC canonical variates. We calculated the silhouette index, a measure of clustering quality, in each of these 5000 distributions and compared the true 4 cluster silhouette index of 0.28 against them (Fig 6C). The true silhouette index of 0.28 was significant relative to the multivariate Gaussian distributions N=143, p=0.049 (Fig 6C). These clusters had unique patterns of RSFC canonical variate scores (Fig 6D). Cluster one had negative and therefore below average RSFC canonical variate values for the global symptom dimension, indicating patients in this group had worse than average symptoms overall. Cluster two had below average RSFC canonical variate scores for the pain catastrophizing dimension, indicating patients in this group had above average catastrophizing. Cluster four had below average pain/anxiety inverse dimension scores, indicating patients in this group had below average mean pain and above average anxiety. Cluster three had above average scores in all three dimensions, indicating patients in this group had the opposite pattern of symptoms as all other biotypes.

## Discussion

Migraine is a heterogeneous disorder with variable response to therapy and expression of symptoms. This variability has been ignored in many studies that treat migraineurs as a single group in between-subjects analyses contrasting patients and healthy controls. When variability has been factored in, subjects are often subtyped into categories not derived from biological data that have also not been rigorously tested. When studies do perform within-subjects analyses to associate symptoms with biological data, they often examine only one variable of interest, ignoring the complex relationship between migraine symptoms. In the current study we addressed these existing limitations by using a large sample size, including a wealth of clinical data from well phenotyped subjects, and by relating these clinical profiles to RSFC in a single multivariate statistical step, as opposed to a standard mass univariate analysis. This approach identified three dimensions of covariation between clinical and RSFC data. The first was a global-symptom dimension that was associated with all symptoms and was primarily related to frontoparietal network (FPN) and dorsal attention network (DAN) functional connectivity with other networks. The FPN includes posterior parietal cortex and lateral prefrontal cortex and is consistently associated with cognitive control (Bush & Shin, 2006; Scolari et al., 2015) as well as acute and chronic pain (Seminowicz et al., 2011; Seminowicz & Moayedi, 2017; Y. Wang et al., 2017). The DAN contains the intraparietal sulcus and frontal eye fields and is consistently involved in attention (Vossel et al., 2014). Our findings suggest that cognitive networks relate to symptoms generally, and that they may function as targets to simultaneously treat all symptoms at once. Consistent with this, the normalization of FPN functional connectivity has been associated with clinical improvement in migraine (Li et al., 2015). The second dimension reflected inverse pain and anxiety scores and was related to default mode (DMN) and sensorimotor network connectivity. The DMN contains angular gyrus, posterior cingulate cortex, and medial prefrontal cortex and the sensorimotor network contains pre- and postcentral gyri. Finally, the third dimension was a pain catastrophizing dimension and was primarily associated with functional connectivity of salience network (anterior insula and midcingulate cortex) to DMN and sensorimotor networks, and to DMN connectivity with other networks.

We were able to show that the RSFC scores for the global symptom dimension was associated with improvements from the non-pharmacologic MBSR+ treatment of migraine, which has recently been established efficacy for migraine (Seminowicz et al., 2020). Subjects with lower scores and thus more severe symptoms showed a larger reduction in headache frequency following MBSR+ than subjects who had comparatively less severe symptoms. In other words, patients with more severe symptoms overall experienced more pronounced improvements from MBSR+. Additionally, we observed that changes in global dimension RSFC scores towards the positive pole (less symptom severe) were associated with improvements from MBSR+ as well. This might indicate that reducing FPN connectivity to salience, sensorimotor, and dorsal attention networks could enhance the effects of MBSR+, given that the first dimension was negatively associated with this FPN functional connectivity.

The migraine RSFC literature uses small sample sizes and inconsistent analyses, making it difficult to synthesize (Skorobogatykh et al., 2019). However, there do appear to be important points of convergence between our findings and previous work, which further supports the validity of our findings. Abnormal connectivity in migraine is often observed in the FPN, salience, sensorimotor, DMN, and DAN (Hubbard et al., 2014; Jin et al., 2013; Ke et al., 2020; Russo et al., 2012; Xue et al., 2012). We therefore anticipated that these networks would be associated with illness characteristics. Two clinical variables included in our analysis that have been the subject of multiple studies in migraine are headache frequency and pain catastrophizing. Frequency was most related to our first identified global dimension, and associations between headache frequency and functional connectivity of salience and FPN nodes (Hubbard et al., 2014; Mainero et al., 2011; Maleki et al., 2012). Pain catastrophizing is often linked to attentional processes (Quartana et al., 2009) and PCS scores are related to anterior insula (key node of the salience network) and sensorimotor connectivity in migraine patients (Hubbard et al., 2014).

Our study has major implications to migraine research and clinical trial design. Migraine is currently split into episodic and chronic classifications based on a 15 headache a month cutoff (International Headache Society, 2018).There have been criticisms of the concept of chronic migraine in the literature (Medrea & Christi, 2018), and in particular over the exact cutoff point between episodic and chronic patients (Torres-Ferrús et al., 2017). We tested the quality of the current two cluster solution for migraine patients in our data based on seven clinical features, finding no evidence against the null hypothesis of a multivariate normal distribution (which does not have clusters). This is especially interesting because the episodic and chronic migraine patient cohorts in our study were collected at two separate sites, which if anything should bias the results toward a chronic versus episodic clustering solution. Additionally, we identified four biotypes of migraine that could be used for predicting therapeutic outcomes in future studies. Patients falling into a given cluster had unique brain-symptom relationships arguing for targeted therapy (Fig 6D). For instance, patients in cluster one had the most extreme global symptom dimension score, which was associated with FPN and DAN connectivity, and perhaps these patients would benefit the most from interventions targeting these networks. On the other hand, cluster two patients who had more extreme pain catastrophizing might benefit the most from interventions that reduce catastrophizing and also target sensorimotor and salience networks. Together, these clustering results argue against the current episodic/chronic clustering that dominates migraine research and clinical trial design, and suggest an alternative based on clinical and biological association.

This study is not without limitations. A major danger when conducting CCA is overfitting, and previously exciting results in depression may have fallen victim to this issue (Dinga et al., 2019). In the current study we took several steps to mitigate overfitting. First, we reduced the number of features in the model through parcellations (instead of voxels) for RSFC, feature selection, and PCA. Altogether, this allowed us to greatly increase the ratio of subjects to model features, attenuating overfitting. We took the additional step of performing regularized CCA and tuned lambda parameters based on the held out strength of the first canonical correlation. This resulted in very minimal overfitting for the first canonical correlation. However, overfitting was larger for the second and third canonical correlations, indicating that some caution is warranted when interpreting and using them. Future work could expand on our results by testing this model in new data and benefit from data sharing to create larger sample sizes that are less prone to overfitting. Additionally, statistical inference for clustering is challenging (Dinga et al., 2019; Liu et al., 2008). In the current study we tested against a null distribution of a multivariate Gaussian distribution, which when true, would not merit clustering. However, even with rejection of this null hypothesis, it is possible that clustering is still unwarranted because not all multivariate non-Gaussian data are necessarily organized into clusters. Future work could examine the utility of this clustering solution by using the four biotypes to predict therapeutic outcomes from migraine treatment, and by examining if these biotypes exist in additional chronic pain disorders.

In conclusion, we identity an association between migraine symptoms – including headache severity, quality of life, affective measures, and coping – and whole-brain resting state functional connectivity, yielding three dimensions of association between these two domains. These results may facilitate the development of personalized medicine, which is limited by treating illnesses as homogeneous groupings. Moreover, the biological association with our clinical data identifies potential targets for therapy and research. Finally, our data argue against the current clustering of migraine patients into chronic and episodic classifications and instead offer an alternative that is grounded in clinical presentation and biology simultaneously.

## Acknowledgements

The authors declare no conflicts of interest.

## Funding

NCCIH/NIH R01 AT007176 to DAS; generous gifts from the Wings Foundation, the Wintercreek Foundation, and the Higgins Family Trust to BS and RC.

## Notes

### Competing Interest Statement

The authors have declared no competing interest.

